# Searching Post-translational Modifications in Cross-linking Mass Spectrometry Data

**DOI:** 10.1101/2024.03.20.585841

**Authors:** Chen Zhou, Shengzhi Lai, Shuaijian Dai, Peize Zhao, Ning Li, Weichuan Yu

**Affiliations:** Department of Electronic and Computer Engineering, The Hong Kong University of Science and Technology, Hong Kong, China; Interdisciplinary Programs Office, The Hong Kong University of Science and Technology, Hong Kong, China; HKUST Shenzhen-Hong Kong Collaborative Innovation Research Institute, Futian, Shenzhen, China; Division of Life Science, The Hong Kong University of Science and Technology, Hong Kong, China

**Keywords:** cross-linking mass spectrometry, post-translational modifications, peptide identification, database search

## Abstract

Cross-linking mass spectrometry (XL-MS) is a technique for investigating protein-protein interactions (PPIs) and protein structures. In the realm of biology, post-translational modifications (PTMs) play a critical role in regulating PPIs and reshaping protein structures. However, the identification of PTMs in XL-MS data poses a great computational challenge and thus remains unexplored. In this study, we introduce SeaPIC, the first XL-MS tool that enables biologists to investigate PTMs in PPIs and protein structures. Our experiments demonstrate the successful identification of PTMs within cross-linked peptides, which were previously undiscovered.

## 1 Introduction

Cross-linking mass spectrometry (XL-MS) ^1–6^ is a technique to study protein-protein interactions (PPIs) and protein structural conformations in a high throughput manner. In contrast to the conventional MS technique ^7^, XL-MS employs a cross-linker to react with protein complexes at the beginning. Cross-linkers are chemical compounds typically consisting of two reaction groups^8–10^. During experiments, these reaction groups can form covalent bonds by binding to specific amino acids within proteins. In a cellular context, when two proteins interact, indicating their spatial proximity, a cross-linker reacts with them, capturing this PPI information. Meanwhile, the crosslinker can also react with amino acids within the same protein but in different spatial positions, enabling the derivation of topological information regarding the protein’s structural conformation. After cross-linkers react with target protein samples, XL-MS follows the traditional MS procedure to generate the first-stage MS (MS1) spectra and second-stage MS (MS2) spectra.

The analysis of MS2 spectra in XL-MS presents a non-trivial computational challenge. Computing cross-link spectrum matches (CSMs) involves considering the combination of any two peptides in the protein sequence database, resulting in a quadratic searching space problem. To tackle this issue, chemists have designed cleavable cross-linkers ^11–14^ and introduced third-stage MS (MS3) spectra, simplifying the problem to a linear peptide task from the wet lab aspect. Tool developers have designed advanced two-step searching methods ^15,16^ and exhaustive searching algorithms ^17^ that can also efficiently handle the task with linear time complexity from the dry lab perspective. However, these advancements only address the fundamental computational challenge in XL-MS data interpretation without considering the occurrence of any post-translational modifications (PTMs).

PTMs^18^ play a pivotal role in regulating biological processes and enabling precise protein interactions. They also impact protein folding, stability, and conformational changes^19^. In XL-MS, the identification of PTMs within cross-linked peptides is of great importance as it sheds light on various biological processes. Studying PTMs in the XL-MS is more intricate and demanding than just identifying the peptide sequences. It offers valuable insights but remains a complex and relatively unexplored area of research.

In this paper, we present a method for <u>sea</u>rching <u>P</u>TMs in <u>C</u>SMs, SeaPIC, the first tool in the XL-MS community that empowers scientists to uncover PTMs in XL-MS data. SeaPIC can handle various types of spectra, including collision-induced dissociation (CID), high-energy collisional dissociation (HCD), and electron-transfer dissociation (ETD) spectra, and is capable of addressing both non-cleavable and cleavable cross-linking scenarios. Moreover, it can be used in combination with all regular XL-MS search engines. SeaPIC serves as a screening method, utilizing tag information ^20^ to determine the partial sequence of cross-linked peptides and solve potential PTMs in the sequence, without the need to decipher the complete pairs of cross-linked peptides. The workflow of SeaPIC encompasses multiple steps: sequence tag graph construction, fuzzy string matching, fast PTM search, score regularization, and the export of PTM information. By employing SeaPIC’s screening capabilities, researchers can extract PTM information from data, which can be then incorporated as a parameter of variable modifications in any regular XL-MS search engine. To validate the effectiveness of SeaPIC, we conducted experiments using simulated, synthetic, and real experimental datasets. The results demonstrate that SeaPIC enables the discovery of a great number of additional PTM-containing cross-linked peptides in published data that were previously overlooked. This highlights the potential of SeaPIC to uncover valuable insights in XL-MS data and expand our understanding of PTMs.

The rest of the paper is organized as follows. Section 2 describes the SeaPIC’s algorithm. Section 3 demonstrates the validity of SeaPIC on simulated, synthetic, and real datasets. Section 4 concludes the paper and discusses the limitations of SeaPIC.

## 2 Methodology

The workflow of SeaPIC is shown in Fig. 1. SeaPIC starts by extracting tags (sequence fragments) from a given experimental MS2 spectrum and constructing a tag graph to identify the paths. Subsequently, fuzzy string matching is applied to determine the potential sequence of the cross-linked peptides using different tags. Next, the pipeline employs a regularized scoring function and a fast PTM search algorithm to identify the optimal PTM(s) for the peptide sequence. SeaPIC exports the normalized results and provides the significance criterion by using a z-score for each PTM’s information. In the end, we can import SeaPIC’s result into the XL-MS search engine to identify the exact PTM information on each cross-linked peptide.

**Figure 1:**
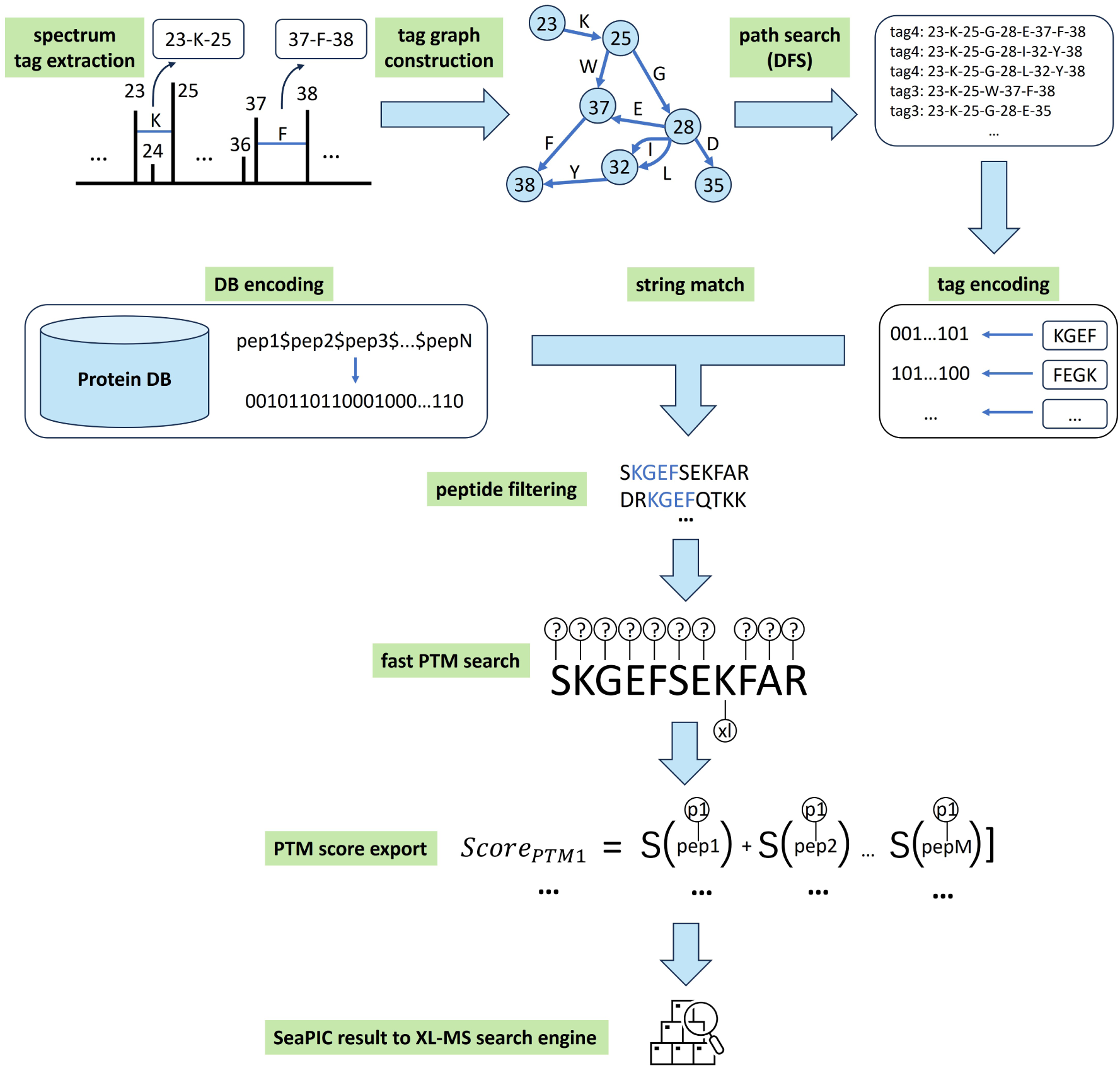
Workflow of SeaPIC. In the query MS2 spectrum, SeaPIC first numbers every peak by their order and pairs every two in a tuple to check if their mass difference is equal to the mass of an amino acid. If so, they will form a tag1, like 23-K-25 and 37-F-38. Then SeaPIC uses all the tag1 information to construct a tag graph, where each vertex represents the peak number and each edge represents the corresponding amino acid. Next, the depth-first search (DFS) algorithm is adopted to find all the paths in the graph, sorting them by path length and average weighted peak intensity. After that, the protein sequence is theoretically digested, and all the peptides are concatenated using the “$” sign. The peptide database and tags are encoded into binary form, and the bitap (shift-or) algorithm is utilized for string matching. The matched peptides are filtered based on the tag position and score. Then, a fast PTM search method is applied to identify possible PTMs on each peptide, finding the best matches using a regularized scoring function. A PTM score is assigned based on the score summation from all the MS2 spectra that identified this PTM, and the results are reported with the z-score significance criterion. Finally, we can import the SeaPIC’s result into a regular XL-MS search engine to find the exact PTM match on the cross-linked peptide.

### 2.1 Tag graph construction

XL-MS data usually has higher charge states and more complex isotopic clusters in MS2 spectra than conventional data. We use the MASCOT Distiller to de-charge MS2 peaks and generate the MGF file. The parameters are shown in our source code (zhouchen xlms ptm.ThermoXcalibur.opt).

In an MS2 spectrum, for every two peaks, we check if their mass difference (delta mass) matches any of the 20 amino acids’ masses within the user-defined error tolerance. If a match is found, we record the locations of these peaks and the corresponding amino acid information as a tuple p1-aa-p2, where p1 represents the location of the lighter mass peak, aa represents the amino acid letter, and p2 represents the location of the heavier mass peak. For instance, 23-K-25 indicates that the 23rd and 25th peaks in the spectra form a lysine mass. This tuple with a length of one amino acid is referred to as tag1. Once the extraction of all tag1 information is completed, we construct a directed graph using these tag1s. In the graph, each vertex represents the location of a peak, and each edge represents the amino acid information.

After constructing the graph, we proceed to identify all the paths within it. To achieve this, we start with vertices that have no incoming edges and treat them as the starting nodes. Using a depth-first search (DFS) approach, we explore all possible paths from these starting nodes to the end nodes in the graph.

Once we have obtained the paths, we prune them and merge similar paths to enhance the robustness of the results. We rank the paths and retain the top N tags in the end. The concrete pruning and filtering procedure can be summarized as follows:

1. Within each path, if a vertex has a significantly lower peak intensity (less than 10%) compared to its neighboring vertices, we remove this vertex. This action breaks the original path into two sub-paths.
2. We compare each pair of paths and check if they have more than 70% overlap. If so, we deduce a new path using the overlap and remove the original two paths. This merging process helps eliminate redundant information and streamline the results.
3. All paths are sorted based on their length and the sum of weighted intensity. The weighted intensity is calculated by dividing the total intensity by the length of the tag. We then retain the top N (default 10) paths for further analysis.
4. Among the top paths, each path should contain at least four amino acids (i.e., at least one tag4), to ensure its significance and relevance.

### 2.2 Fuzzy string match

Given the top N paths derived from the graph, we attempt to find corresponding peptide sequences in the database. Following *in silico* protein digestion (without considering decoy proteins^21^), we combine all the peptide sequences into a single string. We then employ the fuzzy string matching algorithm bitap (also known as shift-or) ^22^ to search for tag patterns (including reverse order for y series ions) in the database, allowing for one amino acid substitution mismatch. The bitap algorithm converts both the concatenated peptide sequence and the tag patterns into binary code and utilizes bit-wise operations to perform string matching. This implementation is highly efficient and exhibits a predictable running time complexity of O(mn), where m represents the length of the concatenated peptide and n represents the length of the pattern.

After performing string matching, we filter the peptides and transform them into potential cross-linked peptides. These transformed peptides then undergo a rough scoring process, and we retain only the highest-ranking results. The procedures are as follows:

1. Peptide filtering.

- The observed peptide mass must not exceed the precursor mass minus the residual mass of the cross-linker.
- If the theoretical peaks for the tag position in the peptide sequence deviate significantly from the corresponding experimental peaks (i.e., outside the mass tolerance range of *±*250 Da), the peptide is considered invalid.
2. Peptide modification.

- To achieve perfect alignment between theoretical peaks and tag peaks in the MS2 spectrum, the mass at the N-terminus of the observed peptide is artificially adjusted for b ions tags (or C-terminus for y ions tags).
- Additional mass is introduced at the link site of the peptide, specifically equal to the difference between the precursor mass and the modified peptide mass obtained in the previous step. This adjustment leads to the formation of a potential cross-linked peptide.
3. Peptide scoring.

- Each modified cross-linked peptide is assigned a score by using the Xcorr scoring function ^23^.
- The top results are retained based on a user-defined threshold.

### 2.3 Regularized score and fast PTM search

We use the Unimod database ^24^ as our reference. It contains about 1,500 PTMs. Our objective is to determine which PTM(s) are present in a given peptide sequence. This task raises a challenge related to the complexity of the search space for PTMs within the peptide sequence. To elaborate, we conduct a numerical evaluation of the total search space for a peptide sequence of length l. We assume that, on average, there could be k distinct PTMs for each amino acid. When considering a single PTM case on the peptide, there are 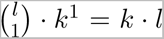 different combinations. For two PTM cases, the number of combinations is 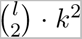, and so on, up to l PTM cases. The total number of PTMs that need to be considered equals to (1 + k)*^l^ −* 1. This term grows exponentially with respect to l and polynomially with respect to k.

The search space is prohibitively large, making it challenging to explore thoroughly. To address this issue, we have developed a fast PTM search method that limits the maximum number of combinations to 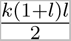. This search space is a polynomial function of l and a linear function of k. Furthermore, we have developed a regularized scoring function S*_regularized_* in Eq. 1:

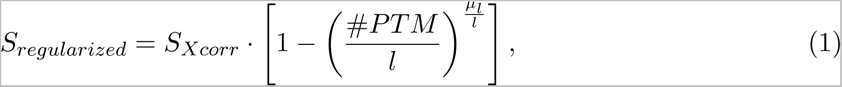

where S*_Xcorr_* represents the original Xcorr scoring function, #PT M denotes the number of PTMs being tested, l denotes the length of peptide, and µ*_l_* represents the average length of peptide candidates for the given experimental spectrum. The rationale behind the scoring function design is elaborated in Supplemental Section 1. This scoring function penalizes the occurrence of PTMs on the peptide sequence. In reality, it is observed that a higher number of PTMs on the peptide corresponds to a lower frequency of occurrence.

In the following, we provide the basic idea and a detailed implementation of the fast PTM search method. For each peptide candidate, we assume it represents one side of the cross-linked peptides and attempt to identify the PTMs on this sequence. If multiple PTMs are present, matching the spectrum to the bald backbone sequence will yield a low score since only a limited number of peaks can be matched. However, as we gradually incorporate the true PTM information into the sequence one by one, the match result should gradually improve. We can continue this process of incrementally adding PTMs until the match results no longer show any significant improvement. By iteratively adding PTMs, we can systematically explore the PTM space and determine the optimal number and type of PTMs in a maximum number of 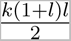 combinations. The pseudo-code is presented in Algorithm. 1.

### Algorithm 1 Fast PTM search algorithm. Inputs are backbone peptide sequence, cross-linking reaction site, PTM map (Unimod database) with amino acids as keys and PTM masses as values, and the precursor mass of cross-linked peptides

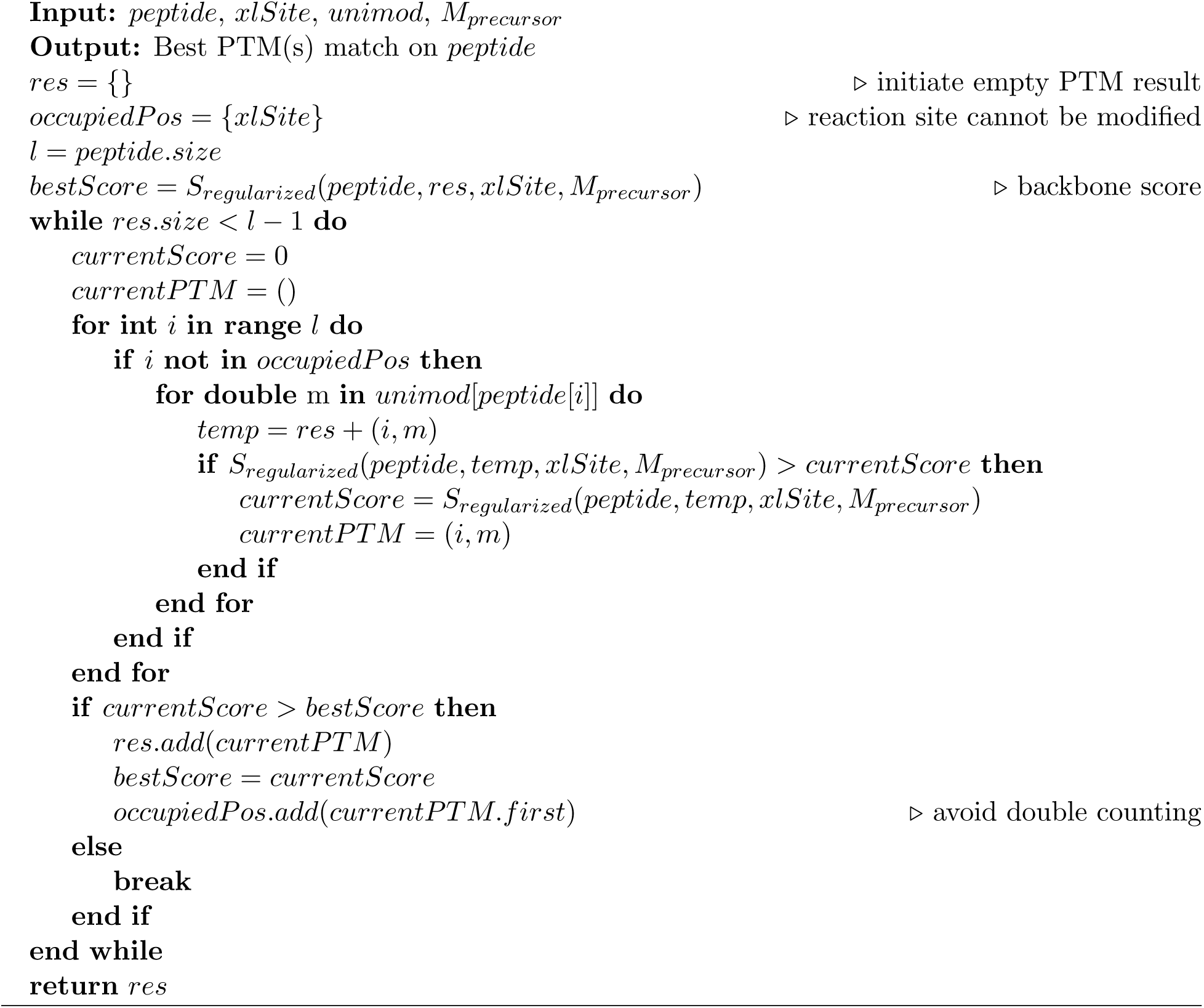

### 2.4 Normalized result

After matching each peptide with the best PTMs using Algorithm 1, the highest-scored peptide along with its corresponding PTMs is considered one of the cross-linked peptides for the current spectrum, unless there are two peptides among the top candidates whose masses exactly add up to the precursor mass. In such cases, these two peptides are treated together as the best matches. Each PTM is then assigned a score in Eq. 2 that aggregates peptide scores across all datasets where this PTM has been identified:

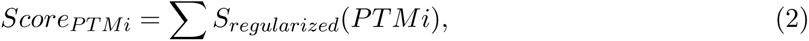

where S*_regularized_*(PT Mi) represents the regularized score for the spectrum that identifies specific

PT Mi.

We have observed that the PTM scores follow a log-normal distribution. By taking the logarithmic form and calculating the z-score for each PTM’s score, we can determine their relative significance. Fig. S3 shows our observation on the distribution of z-scores. We classify the PTM results into four categories based on the following criteria: The absolute value of the lowest z-score is set as the threshold (t). PTMs with z-scores between [1.2t, 1.5t) are considered moderate results, those between [1.5t, 2t) are considered good results, and those between [2t, +*∞*) are considered significant results. The remaining z-scores are labeled as “unsure”.

The PTM information can then be utilized as input for regular XL-MS search engines and serve as a variable modification to search for cross-linked peptides. When determining which PTMs should be included as variable modifications in a regular XL-MS search engine, we recommend considering them in order of high significance to lower significance, including as many as possible while ensuring a suitable running time for the specific search engine.

## 3 Experimental Results

We evaluated the results of SeaPIC through four different experiments, which involved the use of simulated datasets, synthetic isotope-labeled datasets, real experimental datasets, and linear peptide datasets. Throughout the experiments, we tested the SeaPIC in different types of spectra including CID, HCD, and ETD MS2 spectra. The experiments utilized the cleavable cross-linker disuccinimidyl sulfoxide (DSSO) and cyanurbiotindipropionylsuccinimide (CBDPS), and the non-cleavable cross-linker bissulfosuccinimidyl suberate (BS3), and bissulfosuccinimidyl glutarate (BS2G). And we used SeaPIC in conjunction with two different XL-MS search engines, namely pLink2^15^ and ECL3^25,26^ to run out the final PTM-containing CSMs.

### 3.1 Simulated datasets

We initially conducted a simulation experiment to evaluate the performance of SeaPIC. We simulated 100,000 spectra, each with varying numbers and categories of PTMs. We employed SeaPIC to interpret each simulated spectrum, assessing its ability to identify both the backbone sequences and the associated PTMs.

Concretely, we first digested all E.coli proteins into peptides. For each simulation, we randomly selected two peptides from the E.coli database and assumed they were cross-linked peptides. We then introduced several PTMs from the Unimod database onto these cross-linked peptides. The cleavable DSSO cross-linker was used to connect the two peptides. Next, we generated MS2 spectra based on these simulated PTM-containing cross-linked peptides. In reality, the MS2 spectra consist of both fragment (signal) peaks and noise peaks. We assumed that both types of peaks followed a log-normal distribution, with signal peaks having higher intensity compared to noise peaks. Additionally, we assumed that noise peaks were uniformly distributed between 0 Dalton and the precursor mass. For each spectrum, we simulated the presence of 100 noise peaks.

Fig. 2(a) shows an example of the simulated spectrum. We generated a total of 100,000 MS2 spectra and equally divided them into five sets containing 0, 1, 2, 3, and 4 PTMs, respectively. SeaPIC was employed for the analysis of each dataset. While SeaPIC only provides PTM information as output without detailed scan information, we extracted the intermediate outcome from SeaPIC for each spectrum. Subsequently, we compared this intermediate result with the ground truth to determine the sensitivity and precision of the tool. Fig. 2(b)(c) shows the statistics for the backbone sequences and PTMs results. In general, SeaPIC demonstrated a high sensitivity in accurately identifying the majority of backbone sequences and PTMs. It achieves a precision rate of over 90% across various scenarios involving different numbers of PTMs. Fig. 2(b) illustrates the results for backbone sequences. For the spectra set with no PTM, 90.6% of the cross-linked peptides are identified with 99.5% precision. As the number of PTMs increases, the performance of SeaPIC deteriorates, which is reasonable because the PTMs will affect the tag identification. Under the four PTMs situation, SeaPIC still identified 72.1% of the backbone sequences with 97.7% precision. Fig. 2(c) depicts the results of PTM identification. In the spectra set with one PTM, 84.9% of the simulated spectra are identified with 98.6% precision. The detection ability also decreases with an increasing number of PTMs. When there are four PTMs in the spectrum, 62.1% of the results are identified with 92.5% precision. We can observe that even in the four PTMs situation, SeaPIC still identified a big portion of PTMs. We didn’t further simulate more PTM situations in the spectra because we think the case is very rare in reality.

**Figure 2:**
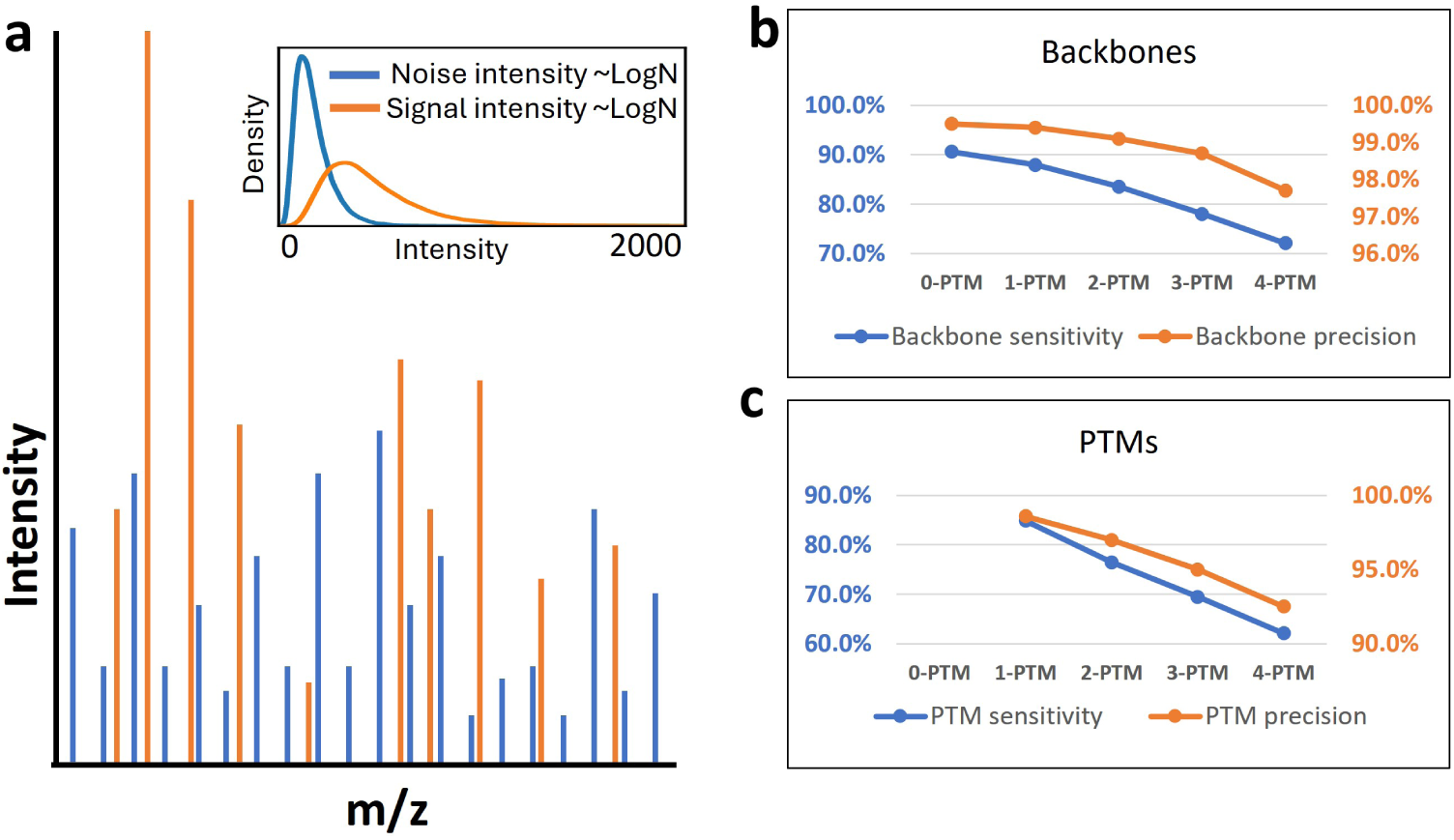
Simulation results. (a) is a demonstration of the simulated spectrum. We randomly selected two peptides from the E.coli database and several PTMs from the Unimod database. Given these artifact peptides, we generated the fragment peaks (b/y ions) with intensity following a log-normal distribution. Additionally, we uniformly generated noise peaks between 0 and the precursor mass, with intensity also following a log-normal distribution but at a lower level. Overall, we simulated 100,000 spectra with 0, 1, 2, 3, and 4 PTMs. (b) shows the performance of SeaPIC in detecting the backbone sequences. When there are no PTMs in the sequence, over 90% of the backbone sequences can be identified with nearly 100% precision. In the case of four PTMs in the peptide sequences, 72% of the backbones are identified with 97% precision. (C) shows the performance of SeaPIC in detecting the PTMs in the sequences. For sequences with one PTM, 85% of the spectra can be identified with nearly 100% precision. When there are four PTMs in the peptide sequences, 62% of the PTMs are identified with 92% precision.

In addition, we employed pLink2 and ECL3 to analyze the aforementioned five datasets with default settings and observe their results. We acknowledge that regular XL-MS search engines struggle to handle PTMs in datasets without prior information. As shown in supplemental Table S1, except for the 0-PTM dataset, both pLink2 and ECL3 failed to identify any CSMs in the remaining datasets. This outcome highlights the importance of SeaPIC in identifying PTMs within XL-MS datasets.

### 3.2 Synthetic datasets

We prepared two sets of cross-linked peptides of bovine serum albumin (BSA) protein using cleavable cross-linker CBDPS^13^. To incorporate PTMs, the peptides were labeled with light (28.0313 Da) and heavy (34.0631 Da) isotope-coded dimethyl chemicals during the sample preparation. Each set of peptides was subjected to a specific acquisition protocol: one set underwent the CID-MS2-ETD-MS2 acquisition protocol, while the other set underwent the HCD-MS2-ETD-MS2 acquisition protocol. The detailed sample preparation procedure is provided in Supplemental Section 2. The mass spectrometry proteomics data have been deposited to the ProteomeXchange Consortium via the PRIDE^27^ partner repository with the dataset identifier PXD048519.

We analyzed the two synthetic datasets using SeaPIC, without specifying the mass of the light and heavy dimethyl. The specific parameters used are included in supplemental Table S2. The results obtained from SeaPIC are presented in Fig. 3. SeaPIC utilizes b/y ions in the CID and HCD spectra, as well as c/z ions in the ETD spectra, to perform the search. The overall results from SeaPIC demonstrate its consistently successful identification of the mass of light and heavy dimethyl. These results exhibit the highest PTM scores and significant confidence. The experiment shows that SeaPIC is capable of fulfilling the task in multiple scenarios, encompassing different spectrum types.

**Figure 3:**
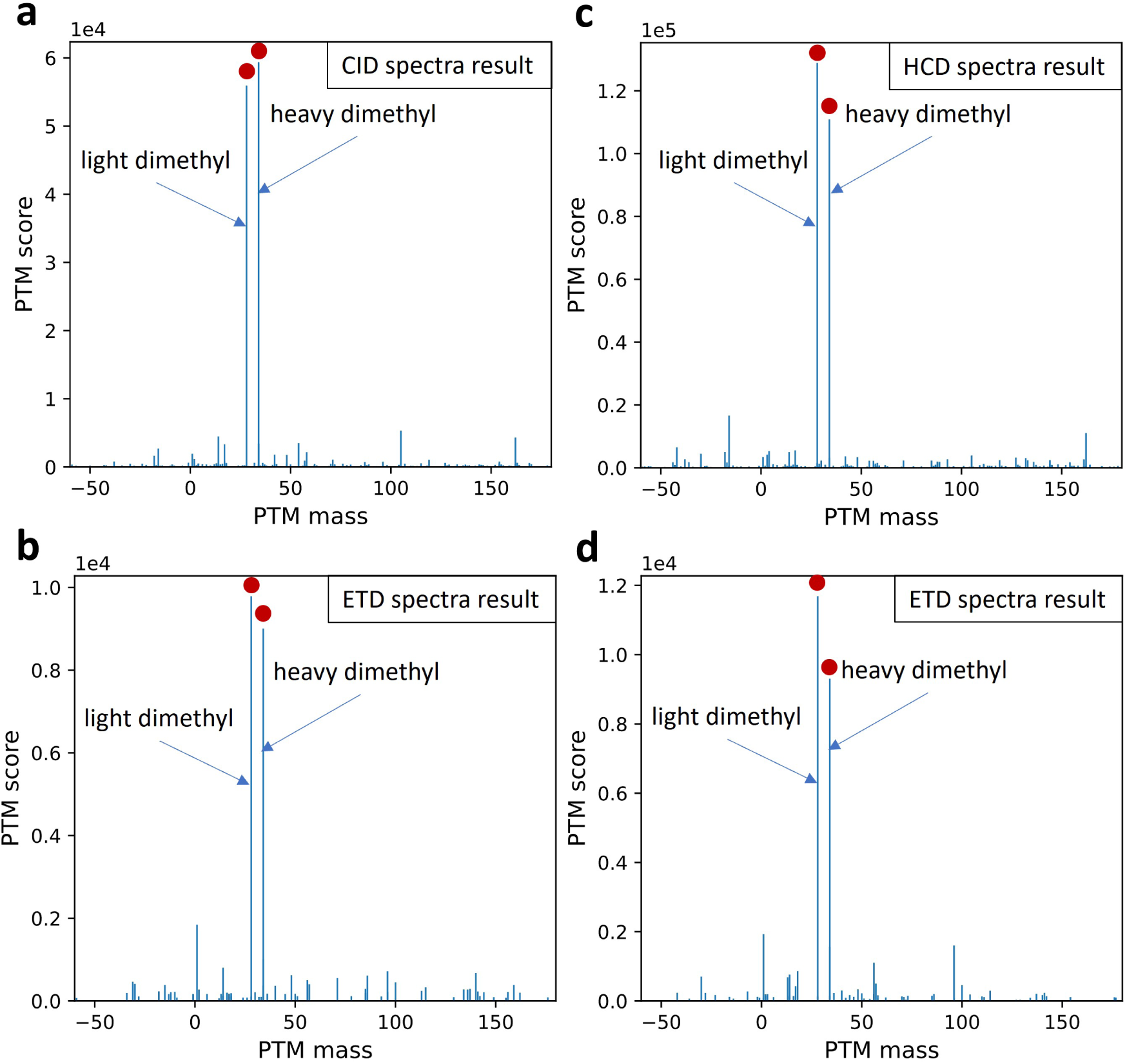
SeaPIC results on synthetic datasets. We utilized SeaPIC to analyze two BSA datasets with known ground truth: CID-MS2-ETD-MS2 and HCD-MS2-ETD-MS2. Bar plots were generated to show the results obtained from SeaPIC, displaying the identified PTM masses along with their corresponding scores. (a) and (b) represent the CID and ETD spectra results from the CID-MS2-ETD-MS2 dataset, while (c) and (d) represent the HCD and ETD spectra results from the HCD-MS2-ETD-MS2 dataset. We can observe that SeaPIC consistently identified the light and heavy dimethyl masses (marked in red circles) in all scenarios, and these identifications were associated with the highest PTM scores. Additionally, the reported z-scores confirmed the significance of these results.

To further compare the effects, we ran pLink2 and ECL3 on these synthetic datasets, both with and without utilizing the PTM information provided by SeaPIC. The parameters used in the search engines are detailed in supplemental Table S3. The number of identified CSMs is summarized in Table 1. It is evident that without the PTM information from SeaPIC, pLink2 and ECL3 yield minimal identifications. However, when provided with the PTM information from SeaPIC, both search engines successfully identified the labeled cross-linked BSA peptides. This contrast can be attributed to the high efficiency of the labeling methods, resulting in the majority of cross-linked peptides carrying mass modifications. Consequently, in the absence of prior knowledge of these modifications, the search engines fail to identify any matches.

**Table 1:**
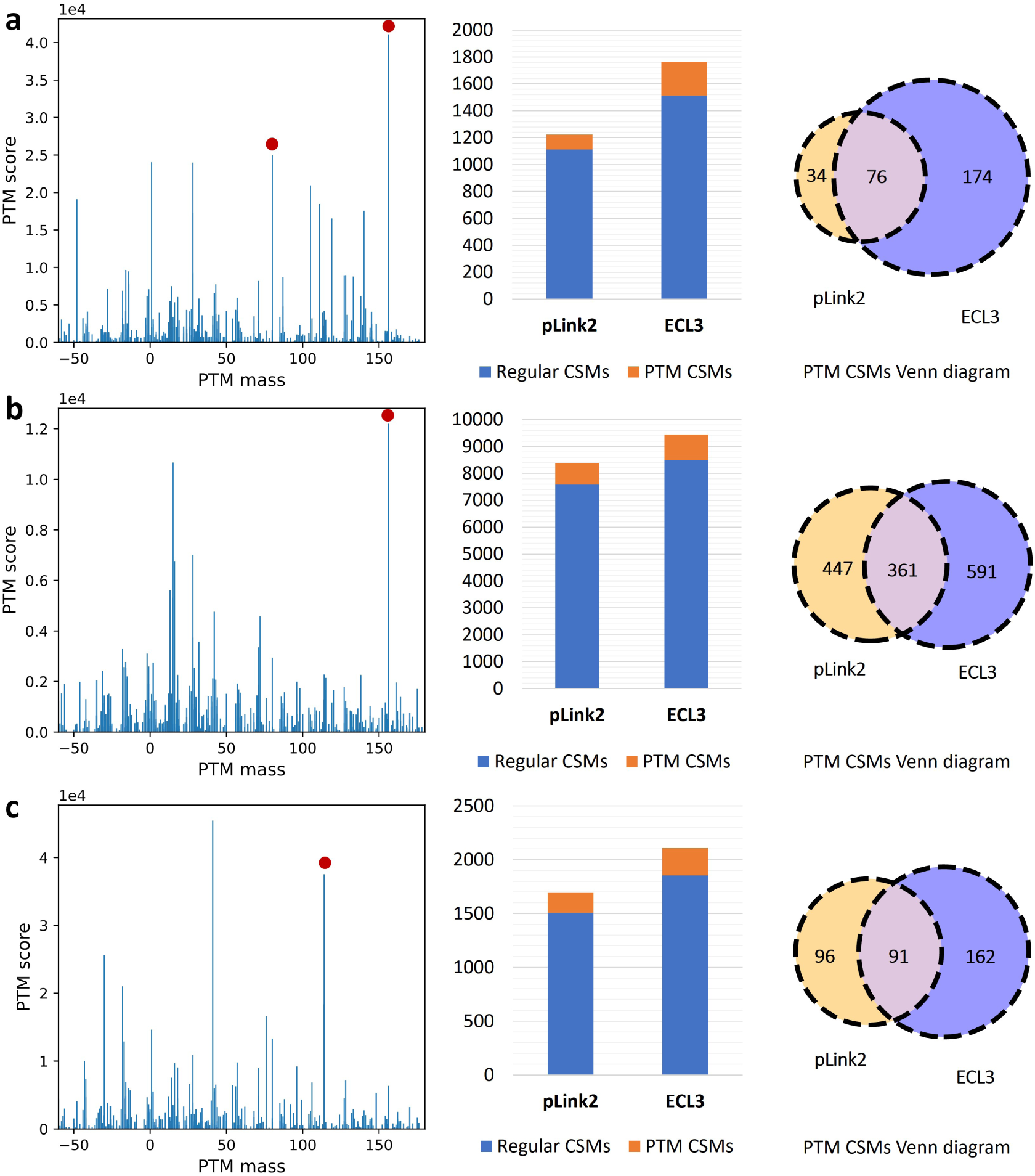
CSMs numbers of pLink2 and ECL3 using/without using SeaPIC on the synthetic dataset. We observed a significant difference in results when we input SeaPIC information into these two XL-MS search engines. This discrepancy can be attributed, in part, to the highly efficient labeling method that leads to the presence of PTMs in almost all cross-linked peptides. Consequently, without prior knowledge of the PTMs in the dataset, the XL-MS search engines struggle to identify any matches.

### 3.3 Real datasets

To investigate the results of SeaPIC on real experimental datasets, we used three human datasets: PXD042584^28^ (BS3 cross-linker), PXD045446^29^ (BS3 cross-linker), and PXD023593^30^ (BS2G cross-linker). We initially analyzed these datasets using SeaPIC. The specific parameters used are included in the supplemental Table S2. Then, we incorporated the PTM information (provided by SeaPIC) into the search engines pLink2 and ECL3 as variable modifications. The output CSMs from these search engines were filtered to retain the highly confident ones. Finally, we drew a Venn diagram to compare the PTM’s consistency from two search engines.

Fig. 4(a), (b), and (c) illustrate the detailed results of the experiments conducted on the three datasets. Bar plots of SeaPIC’s output were generated first, with specific masses annotated. These PTM masses or chemical modifications were either provided by the data source or can be derived from cross-linkers (mono-link mass). Hence, we consider them as true results. All these masses showed high PTM scores, indicating the accuracy of SeaPIC.

**Figure 4:**
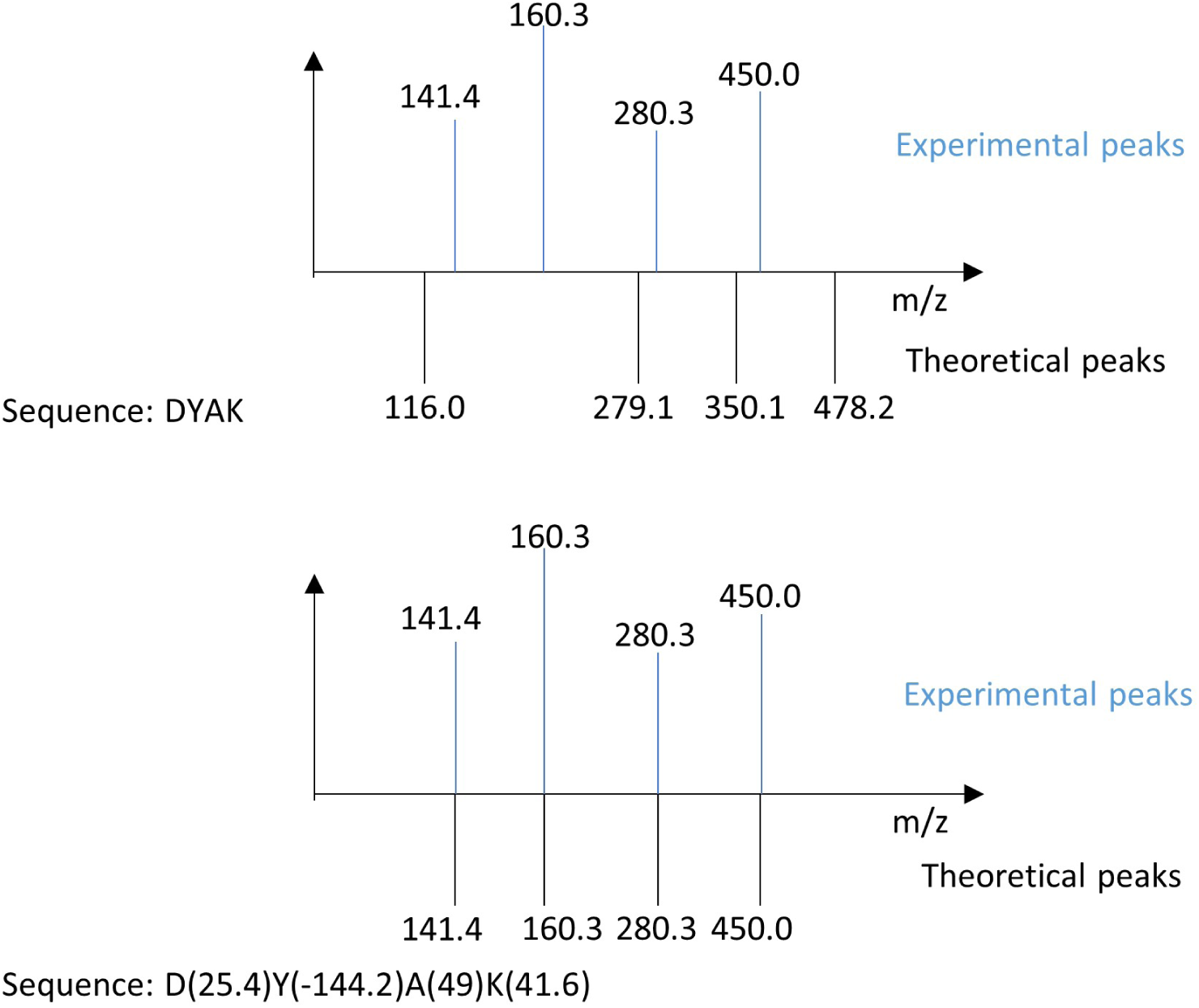
SeaPIC results on experimental datasets. (a), (b), and (c) are the results of PXD042584, PXD045446, and PXD023593, respectively. The datasets underwent the same processing pipeline. Initially, SeaPIC was used to screen the data and generate results on a bar plot. Red circles indicate enriched PTMs or chemical derivatives from a specific cross-linker, that can be verified by the data source. These masses showed high PTM scores. Subsequently, pLink2 and ECL3 were used with the PTM information from SeaPIC. An FDR value of 0.01 was set, and the identified CSMs had to be identified at least twice for output. The CSMs were then classified as regular or PTM, with “regular” referring to results from the original setting and “PTM” indicating results from the SeaPIC setting. The significant number of PTM CSMs (12.4%) suggests that these matches are not random. Finally, a Venn diagram using PTM CSMs was generated. It shows substantial overlap between pLink2 and ECL3, confirming the reliability of the PTM information from SeaPIC.

Next, we used the confident PTM information obtained from SeaPIC as variable modifications in pLink2 and ECL3. The data was then searched, with an FDR threshold of 0.01, and filtered by keeping only redundant CSMs (identified at least twice). The specific searching parameters used for PXD042584, PXD045446, and PXD023593 are included in supplemental Tables S4, S5, and S6. The results were categorized into regular CSMs (identified using original settings) and PTM CSMs (identified using SeaPIC-provided information). We observed that PTM CSMs represented a significant proportion (12.4%) in both pLink2 and ECL3. The FDR threshold, redundant results requirement, and proportion suggest that the presence of PTM-containing CSMs is unlikely to be a random match.

Finally, we examined these PTM CSMs obtained from both pLink2 and ECL3 and visualized the results in a Venn diagram. The overlap area observed further verifies the validity of the PTM information provided by SeaPIC.

### 3.4 Linear peptide datasets

Due to the limited availability of PTM-containing cross-linked peptides with known ground truth, we used an alternative approach to validate SeaPIC. We modified the code to accommodate linear peptide PTM identification, disabling the cross-linking reaction site settings in SeaPIC. This adjustment allowed us to exclusively search for linear peptides with potential PTMs. Furthermore, we had access to numerous publicly available datasets of PTM-containing linear peptides with known ground truth. We conducted tests on 21 different PTMs in linear peptides ^31^ (PXD009449) using SeaPIC and identified all PTMs with the highest PTM scores (supplemental Fig. S4 and S5 and supplemental Tables S7). These experiments demonstrate the reliability of SeaPIC and showcase its applicability in scenarios involving linear peptides. And linear peptide searching mode in SeaPIC shall also benefit the XL-MS datasets because in cleavable cross-linking tasks, MS3 spectra are frequently used to generate the linear peptide data.

## 4 Conclusion

The detection of post-translational modifications (PTMs) in cross-linking mass spectrometry is of great significance as it offers insights into how PTMs impact protein-protein interactions and alter protein structural conformations. Our tool, SeaPIC, is the first of its kind in the community for the investigation of PTMs in cross-linking datasets. SeaPIC has demonstrated its effectiveness by identifying PTM information from publicly available datasets that was previously overlooked. We have also showcased the versatility of SeaPIC in various scenarios, encompassing different spectrum types, cross-linker types, and task types (cross-linked peptides/linear peptides). Nevertheless, SeaPIC has several limitations. In datasets with very sparse PTMs, SeaPIC is capable of identifying them. However, the accumulated PTM score may not effectively differentiate it from the background noise scores. Additionally, it is worth noting that the Unimod database may not contain all naturally occurring PTMs, and SeaPIC is unable to identify PTMs that are not included in the Unimod database.

## 5 Acknowledgements

The work was partially supported by grants 16102422, 16103621, 16306919, 7015-23G, T12-101/23-N, and R4012-18 from Hong Kong Research Grants Council (RGC), grant MHP/033/20 from the Innovation and Technology Commission (ITC) of the Hong Kong S.A.R., the Hetao Shenzhen-Hong Kong Science and Technology Innovation Cooperation Zone project (HZQB-KCZYB-2020083), and internal grants 3030 009 and BGF.001.2023 from HKUST.

## 6 Data and Software Availability

The isotope-labeled BSA dataset can be found on ProteomeXchange under the identifier PXD048519 (reviewer, password: XXXXX). Three human datasets can be found with the identifiers PXD042584, PXD045446, and PXD023593. The PTM-containing linear peptide datasets are available from PXD009449. The SeaPIC program was developed using Python/C++, and can be accessed at https://bioinformatics.hkust.edu.hk or https://github.com/yuweichuan/SeaPIC.git. The running time of SeaPIC varies from several minutes to hours, depending on the sizes of the data and database. Based on our experience using a complete human database (*∼*20,000 proteins) and real datasets (*∼*40,000 MS2 spectra), the screening process typically takes 4-5 hours to complete.

We conducted a concrete running time analysis in supplemental Fig. S6.

## SUPPORTING INFORMATION

### 1 Regularized scoring function

Designing a scoring function for identifying post-translational modifications (PTMs) differs from matching regular peptide sequences. It is essential to consider the probability of PTM occurrence within the peptide sequence. Moreover, it is not always true that a better match of peaks leads to a better result.

When there is no penalty for PTM occurrence within the peptide sequence, we have observed that the program often tends to match PTMs even when they do not exist in reality. This is because adding (fake) PTMs to the sequence will result in a higher chance for the mismatched fragmented ions to match the noisy peaks in the spectrum.

To illustrate the extreme case (Fig. S1), let’s consider a scenario where we have an incorrect backbone sequence that does not have any theoretical ions matching the peaks in the spectrum. However, by deliberately adding PTMs to each amino acid in this backbone sequence, we can generate new theoretical ions that match the peaks in the spectrum. This is possible if there are no limitations on the choice of PTMs, and their mass can be any real number.

Therefore, we need to add a penalty in the scoring function to restrict the occurrence of the PTMs. Concretely, the scoring function needs to satisfy several requirements:

- As the number of PTMs in the backbone sequence increases, the power should also gradually increase.
- If there are no PTMs on the backbone sequence, there shouldn’t be any penalty for the peak matching.
- If all amino acids are assigned PTMs, the score should be 0 regardless of how good the matching result is.
- For peptides with different lengths, the penalty for a single PTM occurrence should vary.

**Figure S1:**
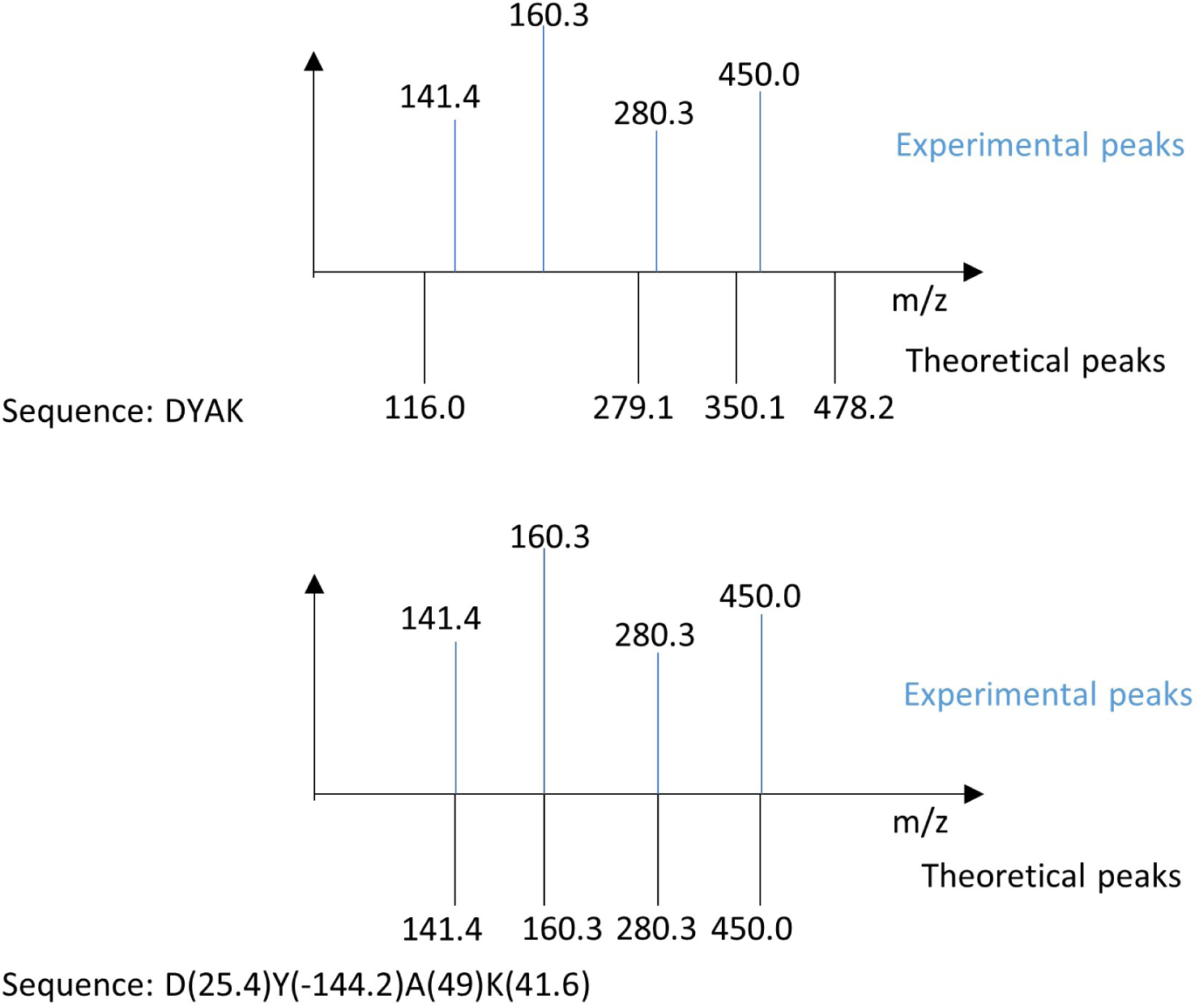
Illustration of the extreme case of fake PTMs on each amino acid to perfectly match the peaks in the spectrum. The sequence ‘DYAK’ cannot match any experimental peaks in the MS2 spectra. But if we add fake PTMs with masses of 25.4Da, -144.2Da, 49Da, and 41.6Da on amino acids ‘D’, ‘Y’, ‘A’, and ‘K’, respectively, we can make the theoretical peaks match the experimental ones perfectly.

Longer peptides should incur a higher penalty. The reason for this is that we can consider a scenario where we have two peptide candidates: A, with a sequence length of 100, and B, with a sequence length of 10. Both sequences A and B have one identified PTM that increases their scores compared to their respective unmodified backbones’ scores. Since A has 100 possible positions to add a PTM and improve the backbone score, while B only has 10 possible positions, we should penalize A more due to its higher random chance of achieving a better score.

Based on the rationales above, we have designed the regularized Xcorr scoring functions^1^:

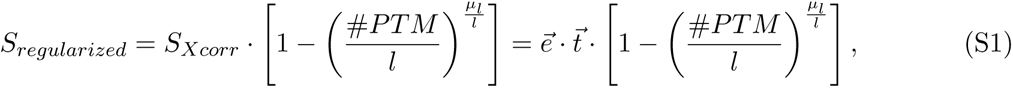

where S*_Xcorr_* represents the original Xcorr scoring function, ⃗e is the digitized experimental peaks, and 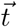 is the digitized theoretical peaks, #PT M denotes the number of PTMs, l denotes the length of peptide, and µ*_l_* represents the average length of peptide candidates for the given spectrum.

To demonstrate the properties of the new scoring function, especially for the weight term 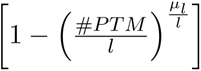 we use some concrete examples and draw a graph.

Suppose we have three peptide candidates for a specific spectrum, with lengths of 5, 10, and 15. Fig. S2 illustrates the curves of the weight changes corresponding to these three lengths. We can observe that the term satisfies our requirements.

**Figure S2:**
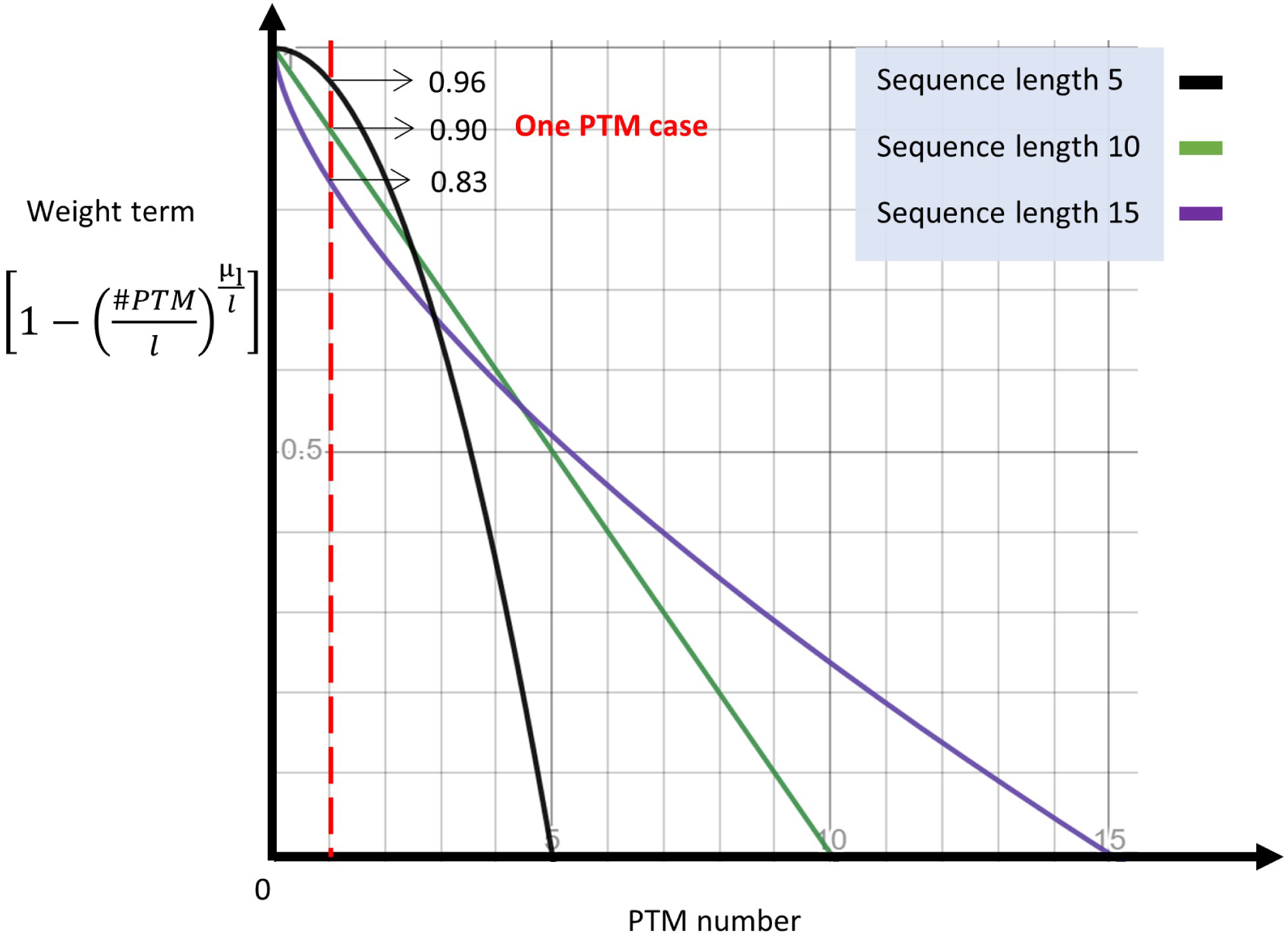
Illustration of the property of the weight term in the scoring function. We generate curves that depict the relationship between the number of PTMs and the weight term in the regularized scoring function. First, all curves are decreasing functions, indicating that as the number of PTMs increases, the weight decreases. Secondly, each curve starts from the point (0,1), indicating that when there are no PTMs (PTM number equals 0), there is no penalty to the score. Third, every curve ends at the point (length, 0), indicating that when all amino acids are assigned PTMs (PTM number equals length), there is a maximum penalty resulting in a weight of 0. Lastly, at the position where the PTM number equals 1, the weight required decreases as the peptide length increases.

### 2 In vitro chemical cross-linking of bovine serum albumin

To prepare the crosslinked bovine Serum Albumin (BSA) protein, 10 mg of BSA was dissolved in 1 mL of crosslinking buffer containing 40 mM HEPES (pH 8.0), 10% glycerol, and 200 mM NaCl. The crosslinker, CBDPS, was freshly prepared in DMSO to achieve a stock solution of 50 mM. The crosslinker was added to the BSA solution at a concentration of 0.5 µM, and the mixture was subjected to 6 rounds of rotation for 3 minutes per round to achieve a final concentration of 3 µM. The mixture was incubated at room temperature for 1 hour, and the excess crosslinker was quenched by adding 1M Tris-HCl (pH 8.0) to the solution to achieve a final concentration of 100 mM, followed by incubation for 10 minutes at room temperature.

The protein pellet was precipitated by applying 3 volumes (v/v) of pre-cooled 12:1 (v/v) acetone/methanol solution for at least 4 hours at -20 °C. The pellet was air-dried and resuspended in protein resuspension buffer 1 containing 50 mM Tris-HCl pH 8.0, 50 mM NaF, 1% glycerol-2-phosphate, 8 M urea, and 2% glycerol. The solution was treated with 10 mM DTT for 30 minutes, 40 mM IAM for 30 minutes (protected from light), and 10 mM DTT for 10 minutes at room temperature. The solution was then mixed with 3 volumes (v/v) pre-cooled 12:1 (v/v) acetone/methanol solution for at least 4 hours at -20 °C. The protein concentration was quantified using a DC protein assay (Bio-Rad, Hercules, CA, USA).

The protein pellet was dissolved in protein re-suspension buffer 2 containing 40 mM Tris-HCl (pH 8.0) and 6 M urea. The protein solution was diluted with a pre-heated (37 °C) trypsin digestion buffer (40 mM Tris-HCl, pH 8.0) to ensure the final concentration of urea was lower than 1 M. The cross-linked peptides were desalted using C18 Sep-Pak cartridges (Waters, Manchester, UK) and concentrated using SpeedVac (Thermo Scientific Inc., Waltham, MA, USA). The XL-peptides were labeled with light (L) and heavy (H) isotope-coded formaldehyde chemicals, mixed, and desalted using C18 Sep-Pak cartridges.

The CBDPS cross-linked peptides were enriched using high-capacity streptavidin agarose resin (Pierce, Rockford, IL, USA) and washed with a washing buffer consisting of 50 mM HEPES (pH 7.5). The cross-linked peptides were eluted with 70% ACN and 0.5% FA for 1 hour, and the elution was performed twice. The peptide samples were desalted using Ziptip (MilliporeSigma, Burlington, MA, USA), followed by SpeedVac.

The cross-linked peptides were analyzed using LC-MS/MS. The peptides were separated by a 120-minute gradient elution at a flow rate of 0.3 µL/min with a Thermo-Dionex Ultimate 3000 HPLC system interfaced with a Thermo Orbitrap Fusion Lumos mass spectrometer. The analytical column was the Acclaim PepMapTM RSLC C18 capillary column (75 µm ID, 150 mm length; Thermo Scientific Inc., Waltham, MA, USA). The MS scans were performed in the range of 300 – 2,000 m/z at a resolution of 120 K, and the MS/MS scans were performed at a resolution of 30,000. The 10 most abundant precursors were subjected to a sequential CID-MS/MS and ETD-MS/MS acquisition protocol.

In the CID-MS/MS experiment, the charge state was set as +4 to +8, and the CID normalized collision energy was 30%. For the ETD-MS/MS experiment, the charge-dependent reaction time was enabled, and the Orbitrap resolution was 30 K. For the HCD-MS/MS experiment, the settings were the same as for the CID-MS/MS except the MS/MS activation type was set as HCD and the collision energy was 26%.

### 3 Tables and figures

**Table S1:** We simulated 100,000 spectra and divided them into five datasets, each containing 0, 1, 2, 3, and 4 PTMs. Both pLink2^2^ and ECL3^3^ were employed to analyze the datasets with default settings, but they failed to identify any results in the remaining datasets except for the 0-PTM dataset. In both pLink2 and ECL3, the identified CSMs in the 1-PTM, 2-PTM, 3-PTM, and 4-PTM datasets are incorrectly matched CSMs.

|  |  | Simulated datasets (each contains 20,000 spectra) |  |  |  |  |
| --- | --- | --- | --- | --- | --- | --- |
|  |  | 0-PTM | 1-PTM | 2-PTM | 3-PTM | 4-PTM |
| Identified CSMs number | pLink2 | 19916 | 51 | 113 | 52 | 20 |
|  | ECL3 | 19556 | 44 | 30 | 21 | 18 |

**Table S2:** SeaPIC parameters used in the synthetic and real datasets.

|  |  |
| --- | --- |
|  | SeaPIC |
| Enzyme | Trypsin |
| Miss_cleavages | 2 |
| Min_length | 5 |
| Known Modifications | C+57.02Da<br>(fixed)<br>M+15.99Da<br>(variable) |
| MS2 tolerance | 0.01Da |
| Linker info | CBDPS $m_{xl} = 509.097\text{Da}$<br>BS2G $m_{xl} = 96.021\text{Da}$<br>BS3 $m_{xl} = 138.068\text{Da}$ |
| Link site | K |

**Table S3:** Search engine parameters used in the synthetic dataset.

| Synthetic dataset | pLink2 | ECL3 |
| --- | --- | --- |
| Enzyme | Trypsin | Trypsin |
| Miss_cleavages | 2 | 2 |
| Min_length | 5 | 5 |
| Modifications | (fixed)<br>C+57.02Da<br>(variable)<br>M+15.99Da<br>n-term,K+28.03Da<br>n-term,K+34.06Da | (fixed)<br>C+57.02Da<br>(variable)<br>M+15.99Da<br>n-term,K+28.03Da<br>n-term,K+34.06Da |
| MS1 tolerance | 10ppm | 10ppm |
| MS2 tolerance | 20ppm | 20ppm |
| Linker info | CBDPS $m_{xl} = 509.097\text{Da}$ | |
| Link site | K | K |
| FDR | 0.01 | 0.01 |

**Table S4:** Search engine parameters used in the PXD042584 dataset.

| PXD042584 | pLink2 | ECL3 |
| --- | --- | --- |
| Enzyme | Trypsin | Trypsin |
| Miss_cleavages | 2 | 2 |
| Min_length | 5 | 5 |
| Modifications | (fixed)<br>C+57.02Da<br>(variable)<br>M+15.99Da<br>STY+79.97Da<br>K+156.08Da | (fixed)<br>C+57.02Da<br>(variable)<br>M+15.99Da<br>STY+79.97Da<br>K+156.08Da |
| MS1 tolerance | 10ppm | 10ppm |
| MS2 tolerance | 0.02Da | 0.02Da |
| Linker info | BS3 $m_{xl} = 138.068\text{Da}$ | |
| Link site | K | K |
| FDR | 0.01 | 0.01 |

**Table S5:** Search engine parameters used in the PXD045446 dataset.

|  |  |  |
| --- | --- | --- |
| PXD045446 | pLink2 | ECL3 |
| Enzyme | Trypsin | Trypsin |
| Miss_cleavages | 2 | 2 |
| Min_length | 5 | 5 |
| Modifications | (fixed)<br>C+57.02Da<br>(variable)<br>M+15.99Da<br>L+15.01Da<br>K+156.08Da | (fixed)<br>C+57.02Da<br>(variable)<br>M+15.99Da<br>L+15.01Da<br>K+156.08Da |
| MS1 tolerance | 10ppm | 10ppm |
| MS2 tolerance | 0.02Da | 0.02Da |
| Linker info | BS3 $m_{xl} = 138.068\text{Da}$ | |
| Link site | K | K |
| FDR | 0.01 | 0.01 |

**Table S6:** Search engine parameters used in the PXD023593 dataset.

|  |  |  |
| --- | --- | --- |
| PXD023593 | pLink2 | ECL3 |
| Enzyme | Trypsin | Trypsin |
| Miss_cleavages | 2 | 2 |
| Min_length | 5 | 5 |
| Modifications | (fixed)<br>C+57.02Da<br>(variable)<br>M+15.99Da<br>S+41.03Da<br>K+114.03Da | (fixed)<br>C+57.02Da<br>(variable)<br>M+15.99Da<br>S+41.03Da<br>K+114.03Da |
| MS1 tolerance | 10ppm | 10ppm |
| MS2 tolerance | 0.02Da | 0.02Da |
| Linker info | BS2G $m_{xl} = 96.021\text{Da}$ | |
| Link site | K | K |
| FDR | 0.01 | 0.01 |

**Figure S3:**
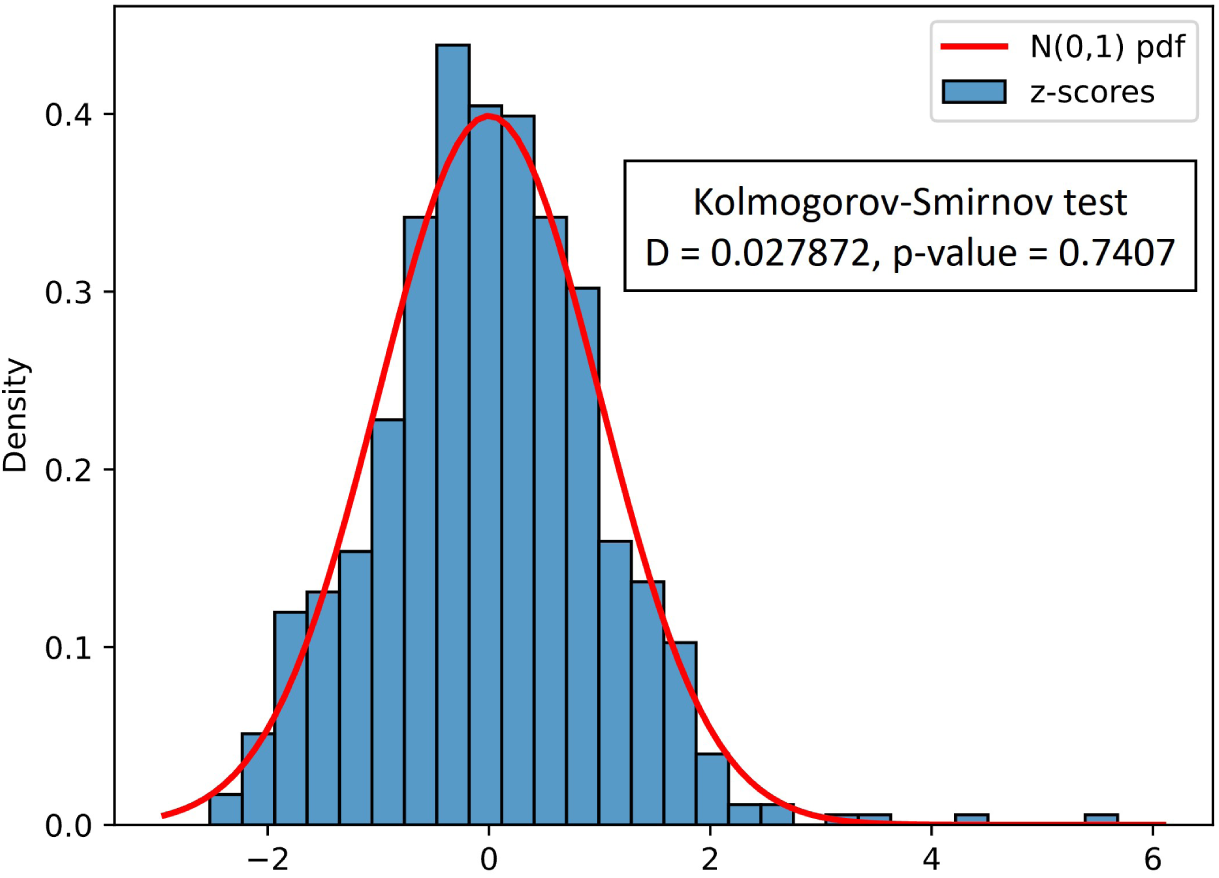
Histogram of z-scores compared with standard normal distribution. We take the logarithmic form of PTM scores and convert them into z-scores by subtracting the mean and dividing by the standard deviation. Then, we plot the histogram of these z-scores and compare it with the standard normal distribution using the K-S test. The p-value indicates that there are no significant differences.

**Figure S4:**
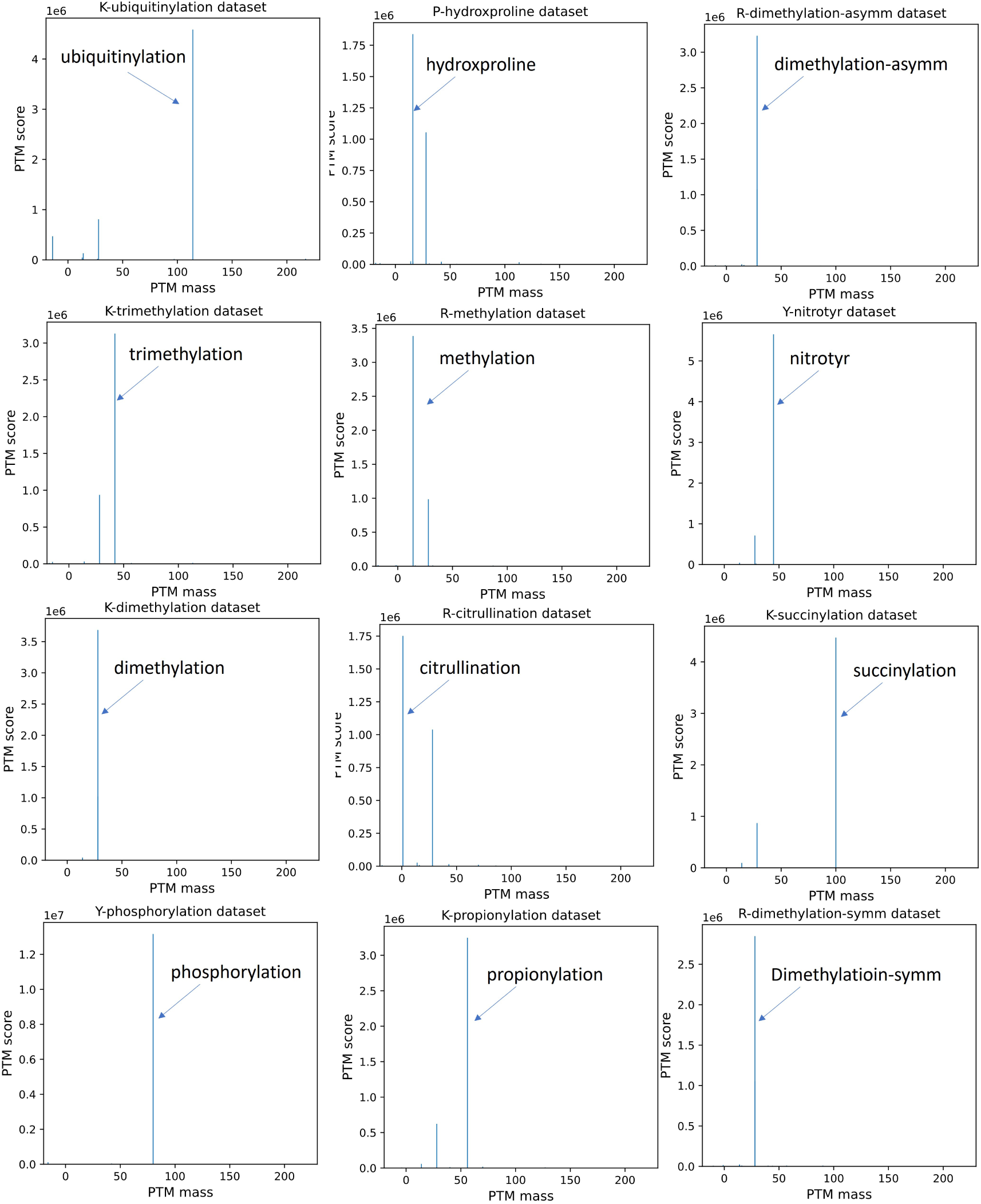
Results of 21 PTMs in the linear datasets. We used SeaPIC to run linear datasets with PTMs and generated bar plots with the horizontal axis PTM mass and the vertical axis PTM score. SeaPIC identified all 21 PTMs as having the highest PTM score.

**Figure S5:**
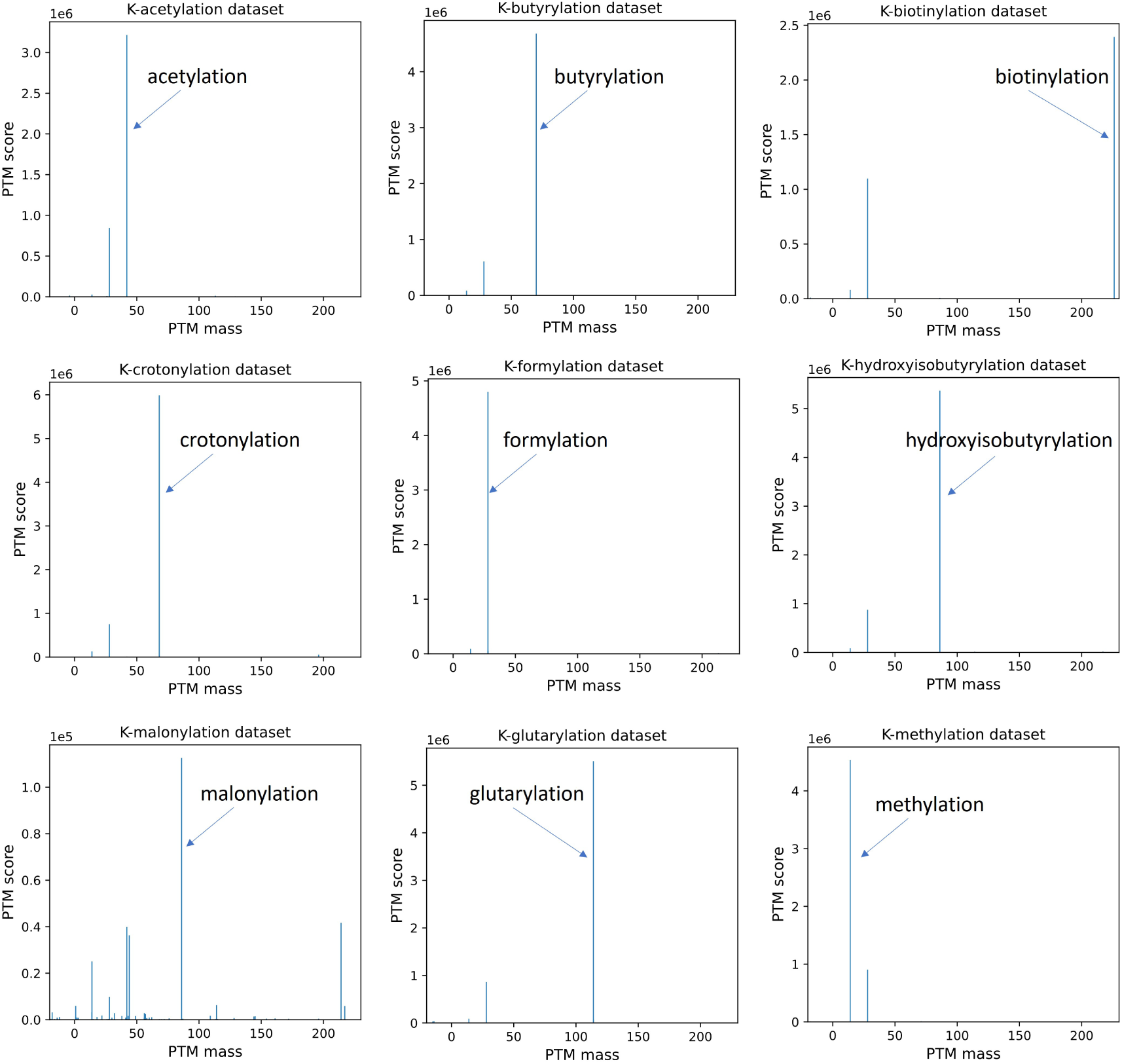
Results of 21 PTMs in the linear datasets. We used SeaPIC to run linear datasets with PTMs and generated bar plots with the horizontal axis PTM mass and the vertical axis PTM score. SeaPIC identified all 21 PTMs as having the highest PTM score.

**Table S7:** Statistics of 21 PTM datasets results for Fig. S4 and Fig. S5. We provided a list of the PTM information and total scan numbers that SeaPIC successfully identified that contained the relevant PTM information.

| file_name | site | PTM name | mass | #scans |
| --- | --- | --- | --- | --- |
| Ymod_Phospho_200fmol_3xHCD_R1 | Y | phosphorylation | 79.966 | 28756 |
| Kmod_Dimethyl_200fmol_3xHCD_R1 | K | dimethylation | 28.031 | 8701 |
| Kmod_Crotonyl_200fmol_3xHCD_R1 | K | crotonylation | 68.026 | 12394 |
| Rmod_Methyl_200fmol_3xHCD_R1 | R | methylation | 14.016 | 8229 |
| Kmod_Trimethyl_200fmol_3xHCD_R1 | K | trimethylation | 42.047 | 7520 |
| Kmod_Hydroxyisobutyryl_200fmol_3xHCD_R1 | K | hydroxyisobutyrylation | 86.037 | 11085 |
| Rmod_Dimethyl_asymm_200fmol_3xHCD_R1 | R | dimethylation-asymm | 28.031 | 7836 |
| Kmod_Formyl_200fmol_3xHCD_R1 | K | formylation | 27.995 | 10488 |
| Kmod_Biotinyl_200fmol_3xHCD_R1 | K | biotinylation | 226.078 | 5323 |
| Rmod_Dimethyl_symm_200fmol_3xHCD_R1 | R | dimethylation-symm | 28.031 | 6967 |
| Kmod_Propionyl_200fmol_3xHCD_R1 | K | propionylation | 56.026 | 6631 |
| Kmod_Ubiquitinyl_200fmol_3xHCD_R1 | K | ubiquitinylation | 114.043 | 9546 |
| Rmod_Citrullin_200fmol_3xHCD_R1 | R | citrullination | 0.984 | 4182 |
| Kmod_Butyryl_200fmol_3xHCD_R2 | K | butyrylation | 70.042 | 10386 |
| Kmod_Succinyl_200fmol_3xHCD_R1 | K | succinylation | 100.016 | 9527 |
| Ymod_Nitrotyr_200fmol_3xHCD_R1 | Y | nitrotyr | 44.985 | 13930 |
| Kmod_Acetyl_200fmol_3xHCD_R1 | K | acetylation | 42.011 | 6685 |
| Kmod_Malonyl_200fmol_3xHCD_R1 | K | malonylation | 86.0 | 10228 |
| Kmod_Methyl_200fmol_3xHCD_R1 | K | methylation | 14.016 | 10216 |
| Kmod_Glutaryl_200fmol_3xHCD_R1 | K | glutarylation | 114.032 | 11553 |
| Pmod_Hydroxyproline_200fmol_3xHCD_R1 | P | hydroxproline | 15.995 | 4143 |

**Figure S6:**
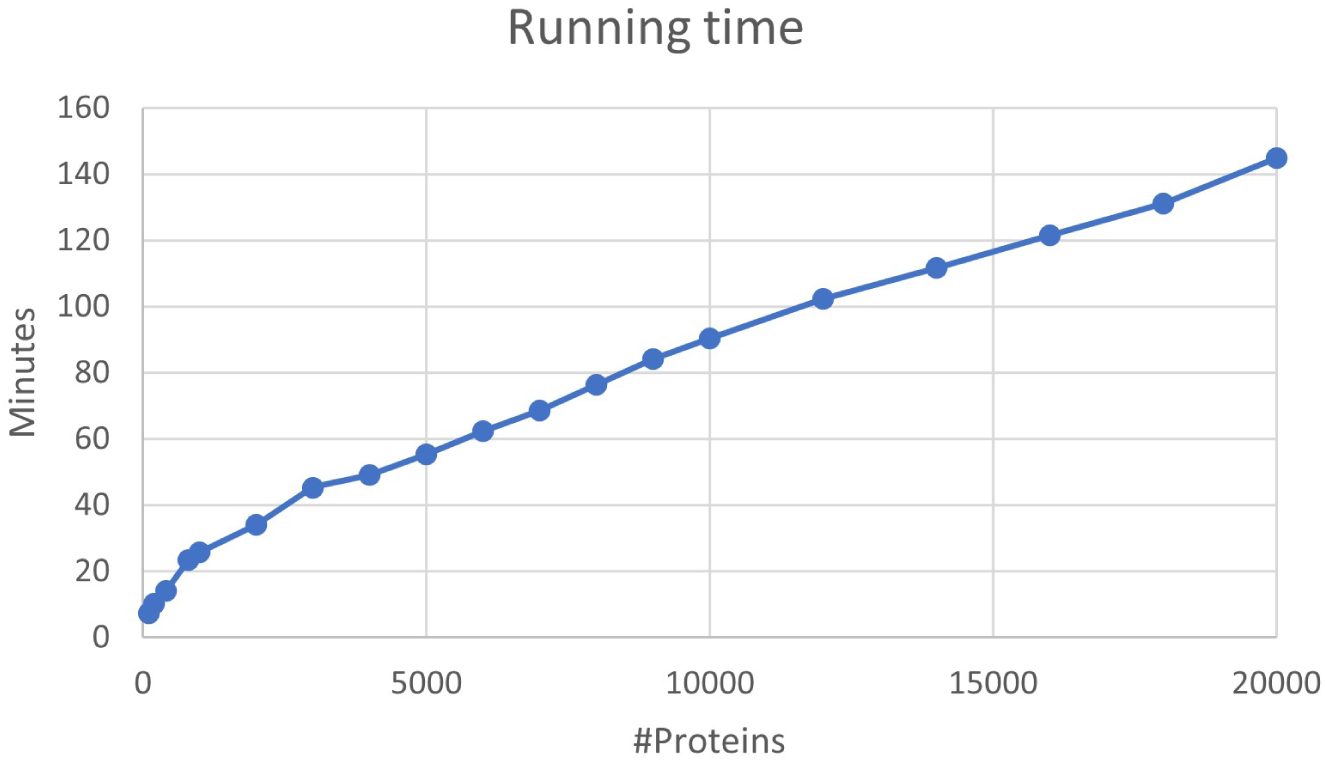
Running time of SeaPIC under different database sizes. SeaPIC runs on an Intel Core i5 2.90 GHz (8 cores/16 processors) Windows desktop computer with 32GB memory. We used different database sizes to run the human dataset with *∼*20,000 spectra. The running time varies from several minutes to hours.

## References

1. A. Leitner, T. Walzthoeni, A. Kahraman, F. Herzog, O. Rinner, M. Beck, and R. Aebersold, “Probing native protein structures by chemical cross-linking, mass spectrometry, and bioinfor-matics,” Molecular & Cellular Proteomics, vol. 9, no. 8, pp. 1634–1649, 2010.

2. A. Sinz, C. Arlt, D. Chorev, and M. Sharon, “Chemical cross-linking and native mass spectrometry: A fruitful combination for structural biology,” Protein Science, vol. 24, no. 8, pp. 1193– 1209, 2015.

3. F. Liu and A. J. Heck, “Interrogating the architecture of protein assemblies and protein interaction networks by cross-linking mass spectrometry,” Current Opinion in Structural Biology, vol. 35, pp. 100–108, 2015.

4. A. Leitner, M. Faini, F. Stengel, and R. Aebersold, “Crosslinking and mass spectrometry: an integrated technology to understand the structure and function of molecular machines,” Trends in Biochemical Sciences, vol. 41, no. 1, pp. 20–32, 2016.

5. C. Iacobucci, C. Piotrowski, R. Aebersold, B. C. Amaral, P. Andrews, K. Bernfur, C. Borchers, N. I. Brodie, J. E. Bruce, Y. Cao, et al., “First community-wide, comparative cross-linking mass spectrometry study,” Analytical Chemistry, vol. 91, no. 11, pp. 6953–6961, 2019.

6. B. Steigenberger, P. Albanese, A. Heck, and R. Scheltema, “To cleave or not to cleave in XL-MS?,” Journal of the American Society for Mass Spectrometry, vol. 31, no. 2, pp. 196–206, 2019.

7. R. Aebersold and M. Mann, “Mass spectrometry-based proteomics,” Nature, vol. 422, no. 6928, pp. 198–207, 2003.

8. C. Clegg and D. Hayes, “Identification of neighbouring proteins in the ribosomes of escherichia coli: A topographical study with the cross-linking reagent dimethyl suberimidate,” European Journal of Biochemistry, vol. 42, no. 1, pp. 21–28, 1974.

9. J. V. Staros, “N-hydroxysulfosuccinimide active esters: Bis (N-hydroxysulfosuccinimide) es-ters of two dicarboxylic acids are hydrophilic, membrane-impermeant, protein cross-linkers,” Biochemistry, vol. 21, no. 17, pp. 3950–3955, 1982.

10. D. Tan, Q. Li, M.-J. Zhang, C. Liu, C. Ma, P. Zhang, Y.-H. Ding, S.-B. Fan, L. Tao, B. Yang, et al., “Trifunctional cross-linker for mapping protein-protein interaction networks and comparing protein conformational states,” Elife, vol. 5, p. e12509, 2016.

11. A. Kao, C.-l. Chiu, D. Vellucci, Y. Yang, V. R. Patel, S. Guan, A. Randall, P. Baldi, S. D. Rychnovsky, and L. Huang, “Development of a novel cross-linking strategy for fast and accurate identification of cross-linked peptides of protein complexes,” Molecular & Cellular Proteomics, vol. 10, no. 1, 2011.

12. A. M. Burke, W. Kandur, E. J. Novitsky, R. M. Kaake, C. Yu, A. Kao, D. Vellucci, L. Huang, and S. D. Rychnovsky, “Synthesis of two new enrichable and MS-cleavable cross-linkers to define protein–protein interactions by mass spectrometry,” Organic & Biomolecular Chemistry, vol. 13, no. 17, pp. 5030–5037, 2015.

13. E. V. Petrotchenko, J. J. Serpa, and C. H. Borchers, “An isotopically coded CID-cleavable biotinylated cross-linker for structural proteomics,” Molecular & Cellular Proteomics, vol. 10, no. 2, pp. S1–S8, 2011.

14. M. Q. Müller, F. Dreiocker, C. H. Ihling, M. Shäfer, and A. Sinz, “Cleavable cross-linker for protein structure analysis: reliable identification of cross-linking products by tandem MS,” Analytical Chemistry, vol. 82, no. 16, pp. 6958–6968, 2010.

15. Z.-L. Chen, J.-M. Meng, Y. Cao, J.-L. Yin, R.-Q. Fang, S.-B. Fan, C. Liu, W.-F. Zeng, Y.- H. Ding, D. Tan, et al., “A high-speed search engine pLink 2 with systematic evaluation for proteome-scale identification of cross-linked peptides,” Nature Communications, vol. 10, no. 1, pp. 3404–3415, 2019.

16. M. R. Hoopmann, A. Zelter, R. S. Johnson, M. Riffle, M. J. MacCoss, T. N. Davis, and R. L. Moritz, “Kojak: efficient analysis of chemically cross-linked protein complexes,” Journal of Proteome Research, vol. 14, no. 5, pp. 2190–2198, 2015.

17. J. Dai, W. Jiang, F. Yu, and W. Yu, “Xolik: Finding cross-linked peptides with maximum paired scores in linear time,” Bioinformatics, vol. 35, no. 2, pp. 251–257, 2019.

18. M. Mann and O. N. Jensen, “Proteomic analysis of post-translational modifications,” Nature Biotechnology, vol. 21, no. 3, pp. 255–261, 2003.

19. O. N. Jensen, “Modification-specific proteomics: characterization of post-translational modifications by mass spectrometry,” Current Opinion in Chemical Biology, vol. 8, no. 1, pp. 33–41, 2004.

20. M. Mann and M. Wilm, “Error-tolerant identification of peptides in sequence databases by peptide sequence tags,” Analytical Chemistry, vol. 66, no. 24, pp. 4390–4399, 1994.

21. J. E. Elias and S. P. Gygi, “Target-decoy search strategy for increased confidence in large-scale protein identifications by mass spectrometry,” Nature Methods, vol. 4, no. 3, pp. 207–214, 2007.

22. R. Baeza-Yates and G. H. Gonnet, “A new approach to text searching,” Communications of the ACM, vol. 35, no. 10, pp. 74–82, 1992.

23. J. K. Eng, B. Fischer, J. Grossmann, and M. J. MacCoss, “A fast sequest cross correlation algorithm,” Journal of Proteome Research, vol. 7, no. 10, pp. 4598–4602, 2008.

24. D. M. Creasy and J. S. Cottrell, “Unimod: Protein modifications for mass spectrometry,” Proteomics, vol. 4, no. 6, pp. 1534–1536, 2004.

25. C. Zhou, S. Dai, S. Lai, Y. Lin, X. Zhang, N. Li, and W. Yu, “ECL 3.0: a sensitive peptide identification tool for cross-linking mass spectrometry data analysis,” BMC Bioinformatics, vol. 24, no. 1, p. 351, 2023.

26. C. Zhou, S. Dai, Y. Lin, S. Lian, X. Fan, N. Li, and W. Yu, “Exhaustive cross-linking search with protein feedback,” Journal of Proteome Research, vol. 22, no. 1, pp. 101–113, 2023.

27. Y. Perez-Riverol, J. Bai, C. Bandla, D. García-Seisdedos, S. Hewapathirana, S. Kamatchinathan, D. J. Kundu, A. Prakash, A. Frericks-Zipper, M. Eisenacher, et al., “The pride database resources in 2022: a hub for mass spectrometry-based proteomics evidences,” Nucleic Acids Research, vol. 50, no. D1, pp. D543–D552, 2022.

28. E. Park, S. Rawson, A. Schmoker, B.-W. Kim, S. Oh, K. Song, H. Jeon, and M. J. Eck, “Cryo-EM structure of a RAS/RAF recruitment complex,” Nature Communications, vol. 14, no. 1, p. 4580, 2023.

29. Y. Chen, G. Kokic, C. Dienemann, O. Dybkov, H. Urlaub, and P. Cramer, “Structure of the transcribing RNA polymerase II–Elongin complex,” Nature Structural & Molecular Biology, pp. 1–11, 2023.

30. C. J. Lupton, C. Bayly-Jones, L. D’Andrea, C. Huang, R. B. Schittenhelm, H. Venugopal, J. C. Whisstock, M. L. Halls, and A. M. Ellisdon, “The cryo-EM structure of the human neurofibromin dimer reveals the molecular basis for neurofibromatosis type 1,” Nature Structural & Molecular Biology, vol. 28, no. 12, pp. 982–988, 2021.

31. D. P. Zolg, M. Wilhelm, T. Schmidt, G. Médard, J. Zerweck, T. Knaute, H. Wenschuh, U. Reimer, K. Schnatbaum, and B. Kuster, “Proteometools: Systematic characterization of 21 post-translational protein modifications by liquid chromatography tandem mass spectrometry (LC-MS/MS) using synthetic peptides,” Molecular & Cellular Proteomics, vol. 17, no. 9, pp. 1850–1863, 2018.

## References

1. J. K. Eng, B. Fischer, J. Grossmann, and M. J. MacCoss, “A fast sequest cross correlation algorithm,” Journal of Proteome Research, vol. 7, no. 10, pp. 4598–4602, 2008.

2. Z.-L. Chen, J.-M. Meng, Y. Cao, J.-L. Yin, R.-Q. Fang, S.-B. Fan, C. Liu, W.-F. Zeng, Y.- H. Ding, D. Tan, et al., “A high-speed search engine pLink 2 with systematic evaluation for proteome-scale identification of cross-linked peptides,” Nature Communications, vol. 10, no. 1, pp. 3404–3415, 2019.

3. C. Zhou, S. Dai, S. Lai, Y. Lin, X. Zhang, N. Li, and W. Yu, “ECL 3.0: a sensitive peptide identification tool for cross-linking mass spectrometry data analysis,” BMC Bioinformatics, vol. 24, no. 1, p. 351, 2023.

